# Mesothelial cell-derived antigen-presenting cancer-associated fibroblasts induce expansion of regulatory T cells in pancreatic cancer

**DOI:** 10.1101/2021.02.04.429827

**Authors:** Huocong Huang, Zhaoning Wang, Yuqing Zhang, Rolf A. Brekken

**Author notes:** **Author Footnotes** Lead Contact (Huocong Huang). **Corresponding authors** Huocong Huang, MD, PhD, Department of Surgery, University of Texas Southwestern Medical Center, 6000 Harry Hines Blvd., Dallas, TX 75390-8593, Rolf A. Brekken, PhD, Department of Surgery and Pharmacology, Hamon Center for Therapeutic Oncology Research, University of Texas Southwestern Medical Center, 6000 Harry Hines Blvd., Dallas, TX 75390-8593.

## Abstract

Multiple recent single cell RNA sequencing (scRNA-seq) studies have identified a unique cancer-associated fibroblast (CAF) population in pancreatic ductal adenocarcinoma (PDA) called antigen-presenting CAFs (apCAFs). apCAFs are characterized by the expression of MHC II molecules, suggesting a function in regulating tumor immunity. Here we integrated multiple scRNA-seq studies and found that apCAFs are derived from mesothelial cells. Our data show that during PDA progression, mesothelial cells form apCAFs by down-regulating mesothelial features and gaining the fibroblastic features, a process induced by IL-1 and TGFβ. Moreover, apCAFs directly ligate and induce naïve CD4^+^ T cells into regulatory T cells (Tregs) in an antigen-specific manner. Our study elucidates an important but neglected cell type in the regulation of PDA immunity and may lead to targeted therapeutic strategies that can overcome immune evasion in PDA.

## Introduction

Pancreatic ductal adenocarcinoma (PDA) is projected to become the second leading cause of cancer-related deaths in the United States by 2030 (Rahib et al., 2014). To date, there is no effective treatment for PDA in part due to the robust desmoplastic reaction that occurs during tumor development (Hosein et al., 2020; Huang and Brekken, 2019). Extensive stroma, which consists of abundant extracellular matrix (ECM) and stromal cells including cancer-associated fibroblasts (CAFs), myeloid cells, lymphocytes and vascular endothelial cells, dominates the PDA microenvironment. CAFs are critical drivers of the desmoplastic reaction, nonetheless, they are also one of the least characterized stromal cells (Chen and Song, 2019; Huang and Brekken, 2020; Kalluri, 2016). Not that long ago, CAFs were considered a group of uniform mesenchymal cells present in the tumor microenvironment (TME) that supported tumor progression. CAFs can promote PDA progression by producing and remodeling ECM and by secreting chemokines, cytokines, and growth factors (LeBleu and Kalluri, 2018). However, ablation studies demonstrated that CAFs can also limit tumor progression (Ozdemir et al., 2014; Rhim et al., 2014). These conflicting data suggest that CAFs consist of heterogeneous subtypes with different functions. CAF heterogeneity was first characterized in an in vitro PDA organoid co-culture model that identified two distinct types of CAFs (Ohlund et al., 2017). One had myofibroblastic features (myCAF), while the other had inflammatory features (iCAF). These two CAF populations were also found to be mutually exclusive and coexist in PDA.

To further understand CAF heterogeneity during PDA progression, we have exploited single-cell RNA sequencing (scRNA-seq) to agnostically profile fibroblasts in three stages of PDA (normal pancreas, early and late PDA) derived from *KIC* (*Kras*^*LSL-G12D*/+^; *Ink4a*^*fl/fl*^; *Ptf1a*^*Cre*/+^) mice (Hosein et al., 2019). We identified three populations of fibroblasts which we named FB1, FB2 and FB3. All three populations were present in normal pancreas and early stage PDA, while only FB1 and FB3 were present in late stage PDA. Among all the fibroblasts populations, FB3 is the most intriguing and unique population as it expresses major histocompatibility complex II (MHC II) genes which are known to be specifically expressed by professional antigen presenting cells. Interestingly, we also found that FB3 is characterized by the expression of mesothelial genes, suggesting that this CAF population may be associated with mesothelial cells. In a later scRNA-seq study in *KPC* (*Kras*^*LSL-G12D*/+^; *Trp53*^*LSL-R172H*/+^; *Ptf1a*^*Cre*/+^) tumors, a CAF population with a similar gene signature as FB3 was identified and termed antigen-presenting CAFs (apCAFs), due to expression of MHC II genes (Elyada et al., 2019). Another scRNA-seq analysis of fibroblasts performed in different stages of *KIC* tumors (normal pancreas, early and established tumor lesions) compared the gene signature of mesothelial cells identified in normal pancreas with apCAFs and found these two cell populations shared similar gene features (Dominguez et al., 2020), further indicating that apCAFs might be derived from mesothelial cells.

Mesothelial cells are derived from embryonic mesoderm (Koopmans and Rinkevich, 2018). During embryonic development, mesothelial cells form a single layer of epithelial cells called coelomic epithelium, which covers the entire coelomic cavity. The coelomic epithelium has multipotency and contributes to growing organs by differentiating into various cell types including fibroblasts and smooth muscle cells through epithelial-to-mesenchymal transition (EMT). The remaining undifferentiated mesothelial cells mature and form a continuous layer of epithelial cells known as mesothelium in the adult. The mesothelium is traditionally thought to be a passive membrane providing a non-adhesive surface covering body cavities, internal organs and tissues. However, recent studies have found that under certain pathological conditions such as wound healing, adipogenesis, myocardial infarction, peritoneal fibrosis and surgical adhesions, mesothelial cells can engage embryonic programs and contribute to the formation of mesenchymal cells (Cao and Poss, 2018; Gupta and Gupta, 2015; Mutsaers et al., 2015; Si et al., 2019; Tsai et al., 2018).

Nonetheless, in cancer, mesothelial cells have not been considered as a functional constituent of the TME. The MHC II-positive CAF populations identified by different scRNA-seq studies underscore that mesothelial cells may have important functions within the TME and may contribute to the formation of a unique CAF population that can regulate tumor immunity by directly interacting with CD4^+^ T cells. Here, we investigate the relationship between mesothelial cells and apCAFs, their functions during PDA progression, as well as the signaling mechanisms that drive the formation of apCAFs.

## Results

### apCAFs are Derived from Mesothelial Cells

In our scRNA-seq study, we found that FB3 exists in multiple genetically engineered mouse models (GEMMs) of PDA regardless of the stage or genotype (normal pancreas, early *KIC*, late *KIC*, late *KPC* and late *KPfC* (*Kras*^*LSL-G12D*/+^; *Trp53*^*fl/fl*^; *Pdx1*^*Cre*/+^)) (Hosein et al., 2019). To understand the relationship of fibroblasts among these models, we projected all the fibroblasts from normal pancreas, early *KIC*, late *KIC*, late *KPC* and late *KPfC* onto a single t-distributed stochastic neighbor embedding (tSNE) plot and applied a graph-based clustering algorithm (Figure 1A). By doing this, we identified four molecular subtypes of fibroblasts (Figure 1B). Two of the four subtypes were what we had identified as FB1, FB2 earlier (Hosein et al., 2019), while the original FB3 cluster further divided into two distinct subtypes, which we named FB3A and FB3B. Interestingly, FB3B expanded specifically in all late stage GEMMs (Figure 1C). We then investigated the transcription profiles of FB3A and FB3B and found that both subtypes were characterized by the expression of MHC II (*Cd74*, *H2-Aa*, *H2-Ab1*, *H2-Dma*, *H2-Dmb1*, *H2-Eb1*) and mesothelial (*Msln*, *Upk3b*, *Ezr*, *Krt19*, *Nkain4*, *Lrrn4*) genes (Figures 1D and 1E). However, in comparison to FB3A, FB3B showed decreased expression of mesothelial genes but an increase of inflammatory (*Il6*, *Cxcl12*, *Pdgfra*) and myofibroblastic (*Acta2*, *Thy1*, *Tgfb1*) genes (Figures 1D and 1F). These data suggest that FB3A and FB3B are mesothelial cells, with FB3A being normal mesothelial cells and FB3B being mesothelial cells with a fibroblastic phenotype.

**Figure 1.**
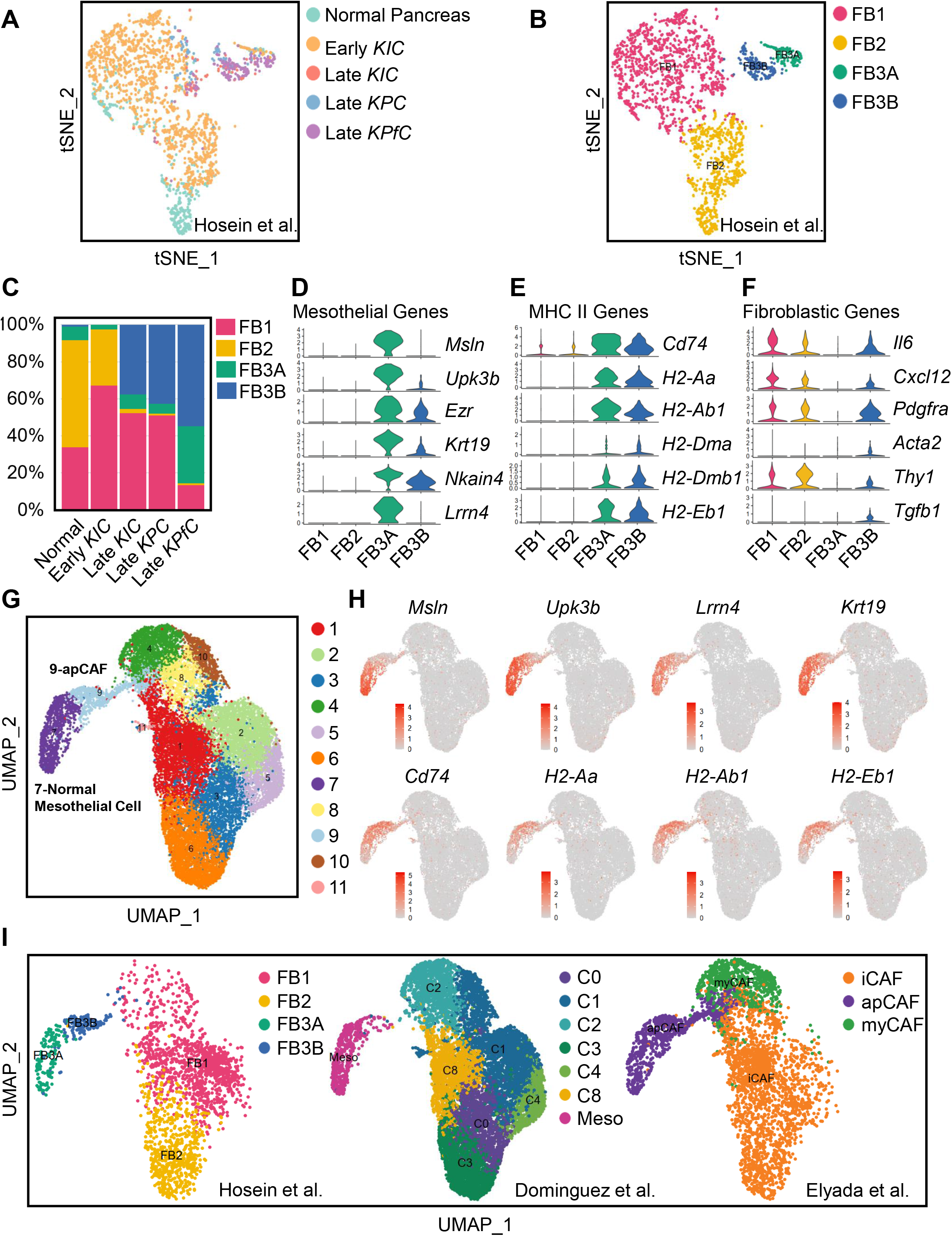
Relationship of Normal Mesothelial Cells and apCAFs in Integrated scRNA-seq Analyses. (A-B) All fibroblasts from scRNA-seq of normal pancreas, early *KIC*, late *KIC*, late *KPC* and late *KPfC* tumor lesions projected onto a tSNE plot with each model (A) or the FB1, FB2, FB3A and FB3B fibroblast populations (B) distinguished by different colors. (C) Proportions of different fibroblast populations in normal pancreas, early *KIC*, late *KIC*, late *KPC* and late *KPfC* tumor lesions, with FB3B specifically expanded in all late stage tumors. (D-F) Violin plots demonstrating marker genes for all the fibroblast populations. Both FB3A and FB3B expressed mesothelial genes (D) and MHC II genes (E), while mesothelial genes were down-regulated (D) and fibroblastic genes were up-regulated (F) in FB3B compared to FB3A. (G) Two other scRNA-seq datasets of fibroblasts in PDA (Dominguez et al., 2020; Elyada et al., 2019) were integrated with our dataset (Hosein et al., 2019). Graph-based clustering of cells with UMAP was performed with the integrated data and 11 clusters of fibroblasts were identified. Further analyses (H-I) suggested that cluster 7 was the normal mesothelial cell cluster and cluster 9 was the fibroblastic mesothelial cell cluster. Here, we defined cluster 9 as apCAFs based on the previous nomenclature of this cell population by Elyada et al (Elyada et al., 2019). (H) Violin plots of mesothelial genes (*Msln*, *Upk3b*, *Lrrn4*, *Krt19*) and MHC II genes (*Cd74*, *H2-Aa*, *H2-Ab1*, *H2-Eb1*) from the integrated fibroblast data. (I) Distribution of fibroblasts in the integrated data based on the origin of datasets (Dominguez et al., 2020; Elyada et al., 2019; Hosein et al., 2019), including FB1, FB2, FB3A and FB3B from Hosein et al. (Hosein et al., 2019), C0-C8 and normal mesothelial cell from Dominguez et al. (Dominguez et al., 2020) and iCAF, apCAF and myCAF from Elyada et al. (Elyada et al., 2019).

Three scRNA-seq studies including ours have identified mesothelial cell-associated cell populations in PDA (FB3A and FB3B from our group (Hosein et al., 2019), apCAF from Elyada et al. (Elyada et al., 2019), normal mesothelial cells from Dominguez et al. (Dominguez et al., 2020)). To understand the relationship of these mesothelial cell-associated populations (Figures 1A and S1A-S1D), we integrated the three fibroblast datasets and performed graph-based clustering of cells with uniform manifold approximation and projection (UMAP). Fibroblasts from the three datasets fell into 11 distinct clusters (Figures 1G and S2). To identify the mesothelial clusters, we generated UMAP plots for MHC II (*Cd74*, *H2-Aa*, *H2-Ab1*, *H2-Eb1*) and mesothelial (*Msln*, *Upk3b*, *Lrrn4*, *Krt19*) genes and found clusters 7 and 9 expressed these signature genes (Figure 1H). We highlighted the distribution of fibroblasts based on the origin of datasets to understand the relationship of FB3A, FB3B, apCAF and normal mesothelial cells (Figure 1I). We found that cluster 7 in the merged data (Figure 1G) had the same features as FB3A while cluster 9 was the same as FB3B, suggesting that cluster 7 represents normal mesothelial cells and cluster 9 represents the fibroblastic mesothelial cell population. Consistently, the mesothelial cell population identified from normal pancreas by Dominguez et al. (Dominguez et al., 2020) overlapped with cluster 7. In addition, we also found that what Elyada et al. (Elyada et al., 2019) termed as apCAFs consisted of normal mesothelial cells and fibroblastic mesothelial cells. Given that cluster 9 was the population that had CAF features, we defined cluster 9 as apCAFs and cluster 7 as normal mesothelial cells in this study (Figure 1G).

### Mesothelial Cells Expand and Contribute to Desmoplasia during PDA Progression

To validate the transcriptomic findings, we performed immunohistochemical (IHC) staining with markers that were mesothelial cell specific. However, we noticed that most of the mesothelial markers such as mesothelin were down-regulated during mesothelial cell to apCAF transition (Figure 2A). Nonetheless, we identified some markers that were specific for mesothelial cells and fibroblasts and were stable during mesothelial cell to apCAF transition (Figure 2A). One of the markers was podoplanin, which was reported by Elyada et al. as a pan-CAF marker (Elyada et al., 2019). Another marker was cadherin-11. IHC for podoplanin, cadherin-11 and mesothelin in normal mouse pancreatic tissue, early and late *KPfC* tumors (Figure 2B) revealed that in normal pancreata, the mesothelium was marked by all three proteins. In early tumor lesions, podoplanin and cadherin-11 marked the expanding mesothelial cells that contributed to the desmoplastic reaction in the niche of early tumor lesions (Figure 2B, arrows). In comparison, mesothelin failed to mark the delaminating mesothelial cells in early PDA lesions. Furthermore, we found that in late stage PDA, the mesothelial layer remained podoplanin and cadherin-11 positive, while mesothelin was down-regulated (Figure 2B). To investigate whether mesothelial cells also contributed to stroma formation in other tumor settings, we established an orthotopic model of PDA using a primary mouse PDA cell line derived from a *KPfC* tumor. Mouse pancreata were harvested one week after implantation, which captured early tumor lesions (Figure 2C). IHC for cadherin 11 revealed that in contrast to the cadherin-11^+^ mesothelium around the uninvolved normal pancreas (Figure 2C, black arrow), cadherin-11^+^ mesothelium adjacent to the tumor expanded and contributed to the tumor stroma (Figure 2C, red arrow). Because podoplanin and cadherin-11 also mark other fibroblast populations, we performed multiplex IHC for podoplanin and the MHC II molecule CD74 to mark mesothelial cells in early and late stage *KPfC* tumors. We found that podoplanin^+^ CD74^+^ mesothelial cells formed a thin membrane lining the edge of pancreas (Figure 2D, left, arrow) and these cells contributed to the TME of early tumor lesions. IHC of late stage tumors indicated that mesothelial cells develop into a major population of CAFs within the tumor stroma (Figure 2D, right, arrow).

**Figure 2.**
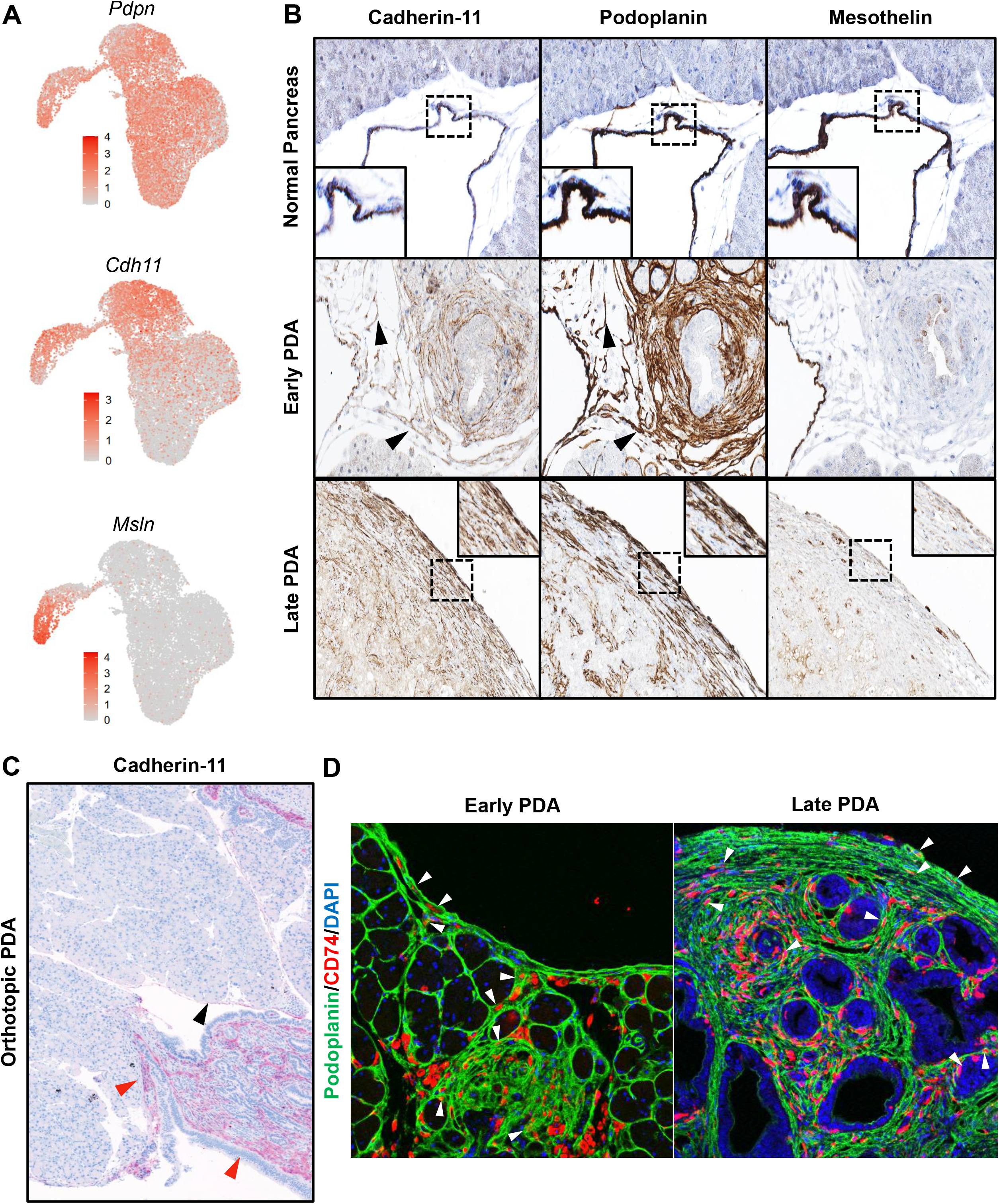
Mesothelial Cells form a Lining of Normal Pancreas and Contribute to the Peripheral CAF formation in PDA. (A) Violin plots of *Pdpn*, *Cdh11* and *Msln* from the integrated fibroblast data. (B) IHC staining of mesothelium in normal pancreata, early *KPfC* tumors (40-day-old) and late *KPfC* tumors (60-day-old) by cadherin-11, podoplanin and mesothelin. Arrows, expanding mesothelial cells. Original magnification 20X. Inset magnification 80X. (C) The primary mouse PDA cell line BMFA3 derived from *KPfC* tumor was orthotopically injected into the pancreata of C57BL/6 mice. Mouse pancreata were harvested one week after the implantation and subjected to IHC staining for cadherin-11. The cadherin-11^+^ mesothelial cells formed mesothelium lining the uninvolved normal pancreas tissues (black arrow). In contrast, the mesothelium adjacent to the tumor lesions started to expand and contributed to the orthotopic tumor stroma formation (red arrow). Magnification 5X. (D) Multiple IHC staining of podoplanin (green), CD74 (red) and DAPI (blue) in early *KPfC* tumors (40-day-old) and late *KPfC* tumors (60-day-old). Arrows, podoplanin^+^CD74^+^ mesothelial cells or apCAFs. Magnification 20X.

To trace the fate of mesothelial cells during PDA progression, we developed a lineage-tracing assay by exploiting a cell-permeable dye, 5(6)-carboxyfluorescein N-hydroxysuccinimidyl ester (CFSE). Once incorporated within cells, CFSE forms stable covalent bonds with intracellular molecules that will be transferred to daughter cells but not adjacent cells (Parish, 1999). This method has been applied to trace the fate of mesothelial cells in a surgical adhesion mouse model (Tsai et al., 2018). We injected CFSE intraperitoneally into wildtype mice and harvested the pancreata after two days. We found that the mesothelium of normal pancreata was labeled by the cell tracking dye (Figure 3A). In contrast, we performed the same assay in 60-day-old tumor-bearing *KIC* or *KPfC* mice and strikingly, we found CFSE-marked cells that infiltrated into the tumor stroma from mesothelial regions (Figure 3A). To ensure the CFSE signal was mesothelium specific, we co-stained the CFSE-labelled tissues with cadherin-11 (Figure 3B). We found that in normal pancreata and PDAs, CFSE^+^ cells were also cadherin-11^+^. In addition, we co-stained with the pancreatic cancer cell marker SOX9 and found that CFSE^+^ cells were distinct from SOX9^+^ cancer cells (Figure 3B), further supporting that the CFSE signal was specific for mesothelium-derived stromal cells.

**Figure 3.**
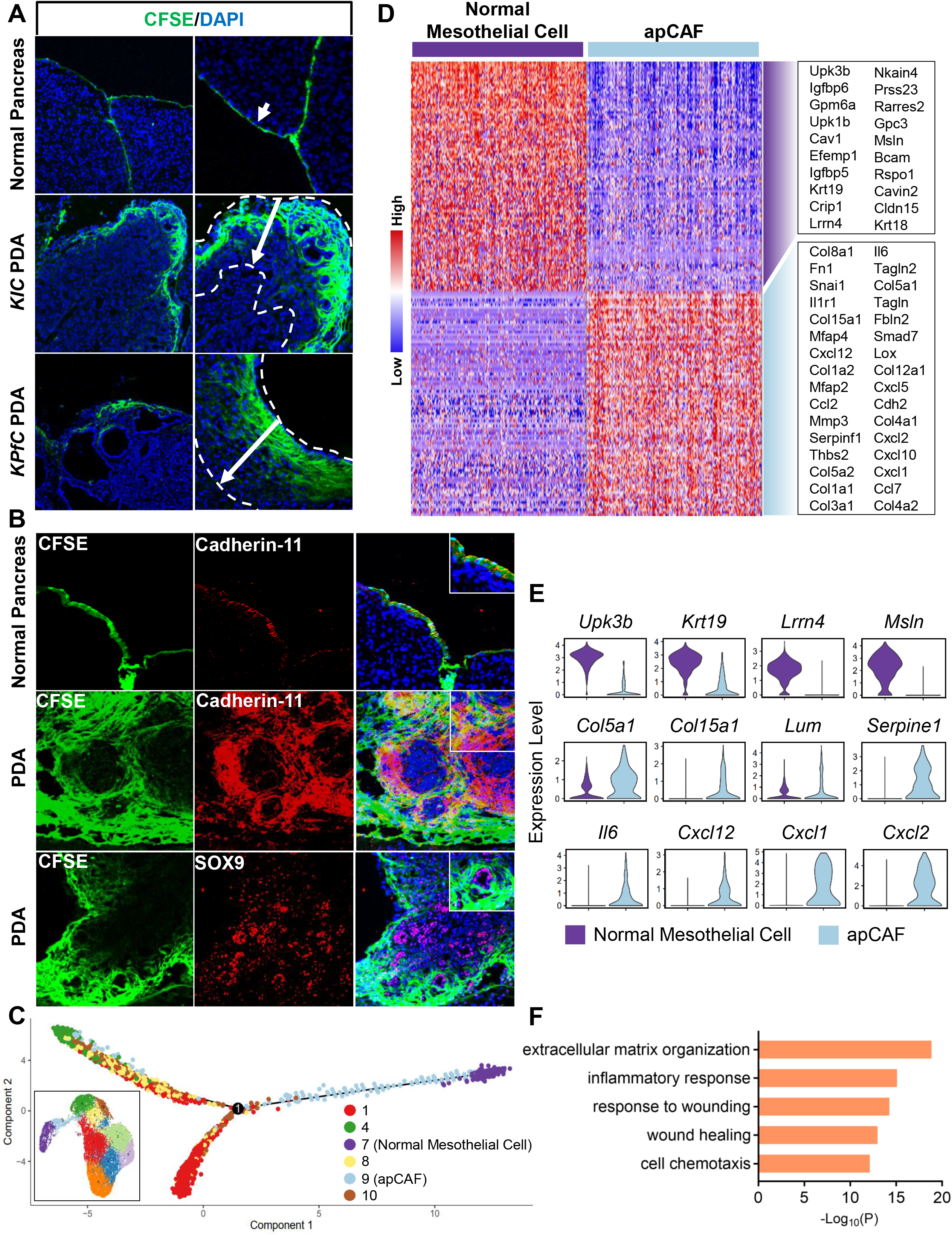
apCAFs are Derived from the Mesothelial Lineage during PDA Formation. (A) Mesothelium tracing with CFSE in normal mice, *KIC* and *KPfC* mice with PDAs (arrows). CFSE was injected intraperitoneally. Normal pancreata or PDAs were harvested 2 days after the injection, then subjected to sectioning and imaging (CFSE, green; DAPI, blue). Magnification 5X (left panel), 10X (right panel). (B) The normal pancreas or *KPfC* PDA tissues from the CFSE lineage tracing assay (A) were stained for cadherin-11 (red) or SOX9 (red) and overlapped with CFSE (green) and DAPI (blue) staining. Original magnification 10X. Inset magnification 20X. (C) Pseudotime analysis with normal mesothelial cells (cluster 7), apCAFs (cluster 9) and other closely associated fibroblast populations (clusters 1, 4, 8, 10) from the integrated data (inset, Figure 1G). (D) The heatmap displaying the top significant genes (cutoff: P<10^−20^) for normal mesothelial cells (cluster 7) and apCAFs (cluster 9) from the integrated data (Figure 1G). (E) Violin plots showing the down-regulation of mesothelial genes (*Upk3b*, *Krt19*, *Lrrn4*, *Msln*) and up-regulation of fibroblastic genes (*Col5a1*, *Col15a1*, *Lum*, *Serpine1*, *Il6*, *Cxcl12*, *Cxcl1*, *Cxcl2*) in apCAFs compared to normal mesothelial cells from the integrated data from the integrated data (Figure 1G). (F) Top biological processes of Gene Ontology (GO) analysis with the up-regulated gene cluster in apCAFs compared with normal mesothelial cells (D).

### Wound-associated Tumor Paracrine Signals Induce the Mesothelial cell-apCAF Transition

In the merged scRNA-seq data of fibroblasts (Figure 1G), the mesothelial cell/apCAF populations had a close relationship with other fibroblast populations (clusters 1, 4, 8, 10). Given that during embryonic development, mesothelial cells can differentiate into fibroblasts and smooth muscle cells (Rinkevich et al., 2012), we performed pseudotime analysis with the normal mesothelial cell (cluster 7), apCAF (cluster 9) and other closely associated fibroblast populations (clusters 1, 4, 8, 10). We found that although normal mesothelial cells gained fibroblastic features and trended towards fibroblasts when they became apCAFs, they had limited contribution to the formation of other CAF populations (Figure 3C), supporting data from Dominguez et al., which suggested that iCAF and myCAF lineages are derived from resident fibroblasts (Dominguez et al., 2020).

Since apCAFs gained a differential gene signature by down-regulating mesothelial genes and up-regulating fibroblastic genes (Figures 3D and 3E), we performed pathway and functional enrichment analysis with the differential up-regulated genes in apCAFs, in an effort to identify the biological processes involved in driving such gene signature changes. We found that many of the identified biological processes were related to wounding or inflammatory response (Figure 3F), suggesting that a wound-associated signal from the tumor niche can induce apCAF formation. We also noticed that the gain of fibroblastic features in mesothelial cells was more obvious in late stage tumors. Although mesothelial cells expanded around the early tumor lesion, these cells did not express fibroblastic markers such as α–smooth muscle actin (α-SMA) (Figure 4A, left). In contrast, the mesothelium in late-stage tumors became fibroblastic (Figure 4A, right). In addition, we co-stained the CFSE^+^ cells of PDA from the lineage-tracing assay (Figure 3A) with α-SMA and found some CFSE^+^ cells were α-SMA^+^ (Figure 4B).

**Figure 4.**
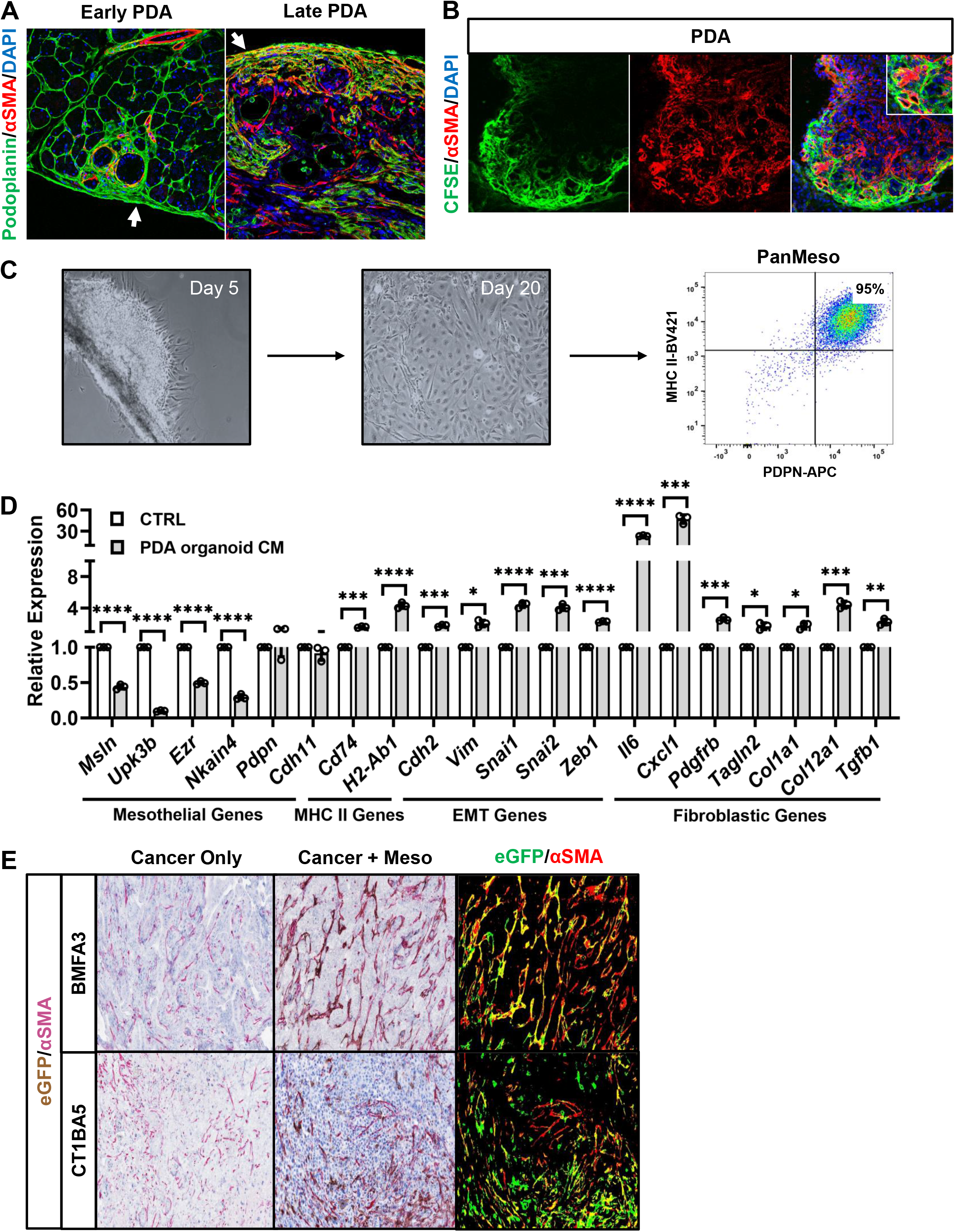
PDA Paracrine Signal Induces Fibroblastic Transition in Mesothelial Cells. (A) Multiple IHC staining of podoplanin (green), α-SMA (red) and DAPI (blue) in early *KPfC* tumors (40-day-old) and late *KPfC* tumors (60-day-old). Arrows, the mesothelium of early or late *KPfC* tumors. Magnification 20X. (B) The *KPfC* PDA tissues from the CFSE lineage tracing assay (Figure 3A) were stained for α-SMA (red) and overlapped with CFSE (green) and DAPI (blue) staining. Original magnification 10X. Inset magnification 20X. (C) The mesothelium tissues from the pancreata of immortomice were isolated, seeded onto a tissue culture dish and cultured at 33 °C. Cells started to migrate from the mesothelium five days after the seeding and became confluent after culturing for 20 days. Cells were subjected to FACS and podoplanin+MHC II+ cells were collected (95%). These cells were named PanMeso cells. (D) The PanMeso cells were treated with PDA organoid conditioned medium derived from *KPfC* tumors for 48 hrs. Cells were harvested and subjected to qPCR for mesothelial (*Msln*, *Upk3b*, *Ezr*, *Nkain4*, *Pdpn*, *Cdh11*), MHC II (*Cd74*, *H2-Ab1*), EMT (*Cdh2*, *Vim*, *Snai1*, *Snai2*, *Zeb1*) and fibroblastic (*Il6*, *Cxcl1*, *Pdgfrb*, *Tagln2*, *Col1a1*, *Col12a1*, *Tgfb1*) genes. n=3, *P<0.05, **P<0.01, ***P<0.001, ****P<0.0001. (E) A primary *KPfC* cancer cell line (BMFA3 or CT1BA5) was injected orthotopically into mouse pancreata with or without PanMeso cells expressing EGFP. Tumors were harvested one month after the injection and stained for the EGFP (brown) and fibroblastic marker α-SMA (red). EGFP was also highlighted as green and α-SMA was highlighted as red by ImageJ (right panel). Yellow marked the EGFP^+^ PanMeso cells expressing fibroblastic marker α-SMA.

To examine if a tumor paracrine signal could induce mesothelial cells to differentiate into apCAFs, we established a mouse pancreatic mesothelial cell line. Mesothelium was harvested from the normal pancreata of immortomice (Jat et al., 1991). Mesothelial explants were seeded onto a tissue culture dish and cultured at 33 °C. Five days after seeding, cells started to migrate from the mesothelium (Figure 4C). Once confluent, these cells were subjected to flow cytometry. Ninety-five % of the cells were podoplanin^+^MHC II^+^ (Figure 4C). They were collected and named pancreatic mesothelial (PanMeso) cells. We treated PanMeso cells with *KPfC* PDA organoid conditioned medium (CM) and performed qPCR (Figure 4D). Treatment with PDA organoid CM significantly down-regulated mesothelial genes (*Msln*, *Upk3b*, *Ezr*, *Nkain4*). Consistent with the scRNA-seq data (Figure 2A), the expression of *Pdpn* and *Cdh11* was not decreased by the CM. Moreover, MHC II genes (*Cd74*, *H2-Ab1*) were up-regulated by the CM. Interestingly, we found that many EMT-related genes (*Cdh2*, *Vim*, *Snai1*, *Snai2*, *Zeb1*) were induced by the CM, suggesting that the embryonic EMT program in mesothelial cells that contributes to the formation of mesenchymal cells in developing organs was reactivated by the tumor paracrine signal (Koopmans and Rinkevich, 2018; Rinkevich et al., 2012). Furthermore, CM induced a fibroblastic transition in the PanMeso cells evidenced by up-regulation of fibroblastic genes including iCAF genes (*Il6*, *Cxcl1*, *Pdgfrb*) and myCAF genes (*Tagln2*, *Col1a1*, *Col12a1*, *Tgfb1*).

To investigate whether the in vivo TME could induce the apCAF phenotype in PanMeso cells, we co-injected eGFP-PanMeso cells with a primary *KPfC* cell line (BMFA3 or CT1BA5). As control, we also injected cancer cells alone. Tumors were harvested one month after injection and subjected to IHC for eGFP and α-SMA (Figure 4E). We found that the TME induced eGFP^+^ PanMeso cells to express the myCAF protein α-SMA^+^. Taken together, the in vitro and in vivo data using PanMeso cells indicate that a tumor paracrine signal can induce a fibroblastic transition in mesothelial cells.

### apCAFs Induce Naïve CD4+ T Cells into Regulatory T Cells

Although apCAFs express MHC II molecules and can present antigen to CD4^+^ T cells, they lack the classical co-stimulatory molecules (e.g., CD40, CD80 and CD86) that are necessary to induce full CD4+ T cell activation and clonal expansion following T-cell receptor (TCR) ligation (Elyada et al., 2019). Therefore, we sought to understand the function of apCAFs in PDA and determine how the interaction between apCAFs and CD4^+^ T cells influences T cell phenotype. First, we compared the antigen-presenting capability of normal mesothelial cells with apCAFs. Podoplanin^+^MHC II^+^ mesothelial cells from normal pancreata or apCAFs from *KPfC* tumors were sorted by FACS (Figure 5A). We incubated the sorted mesothelial cells or apCAFs with the ovalbumin (OVA) peptide (OVA 323-339), and then co-cultured them with CD4^+^ T cells isolated from OT II mice (Barnden et al., 1998). We found that normal mesothelial cells and apCAFs induced T cell expression of early activation markers of TCR ligation, CD25 and CD69, in an OVA-specific manner (Figures 5B and 5D). This effect was more obvious in apCAFs. To validate this result, we performed a similar assay using PanMeso cells (Figure 5A). PanMeso cells were pre-treated with control medium or PDA organoid CM for 2 days before the incubation with OVA peptide. We found that the formation of apCAFs induced by PDA organoid CM led to an increase of antigen-presenting capability in PanMeso cells (Figures 5B and 5D).

**Figure 5.**
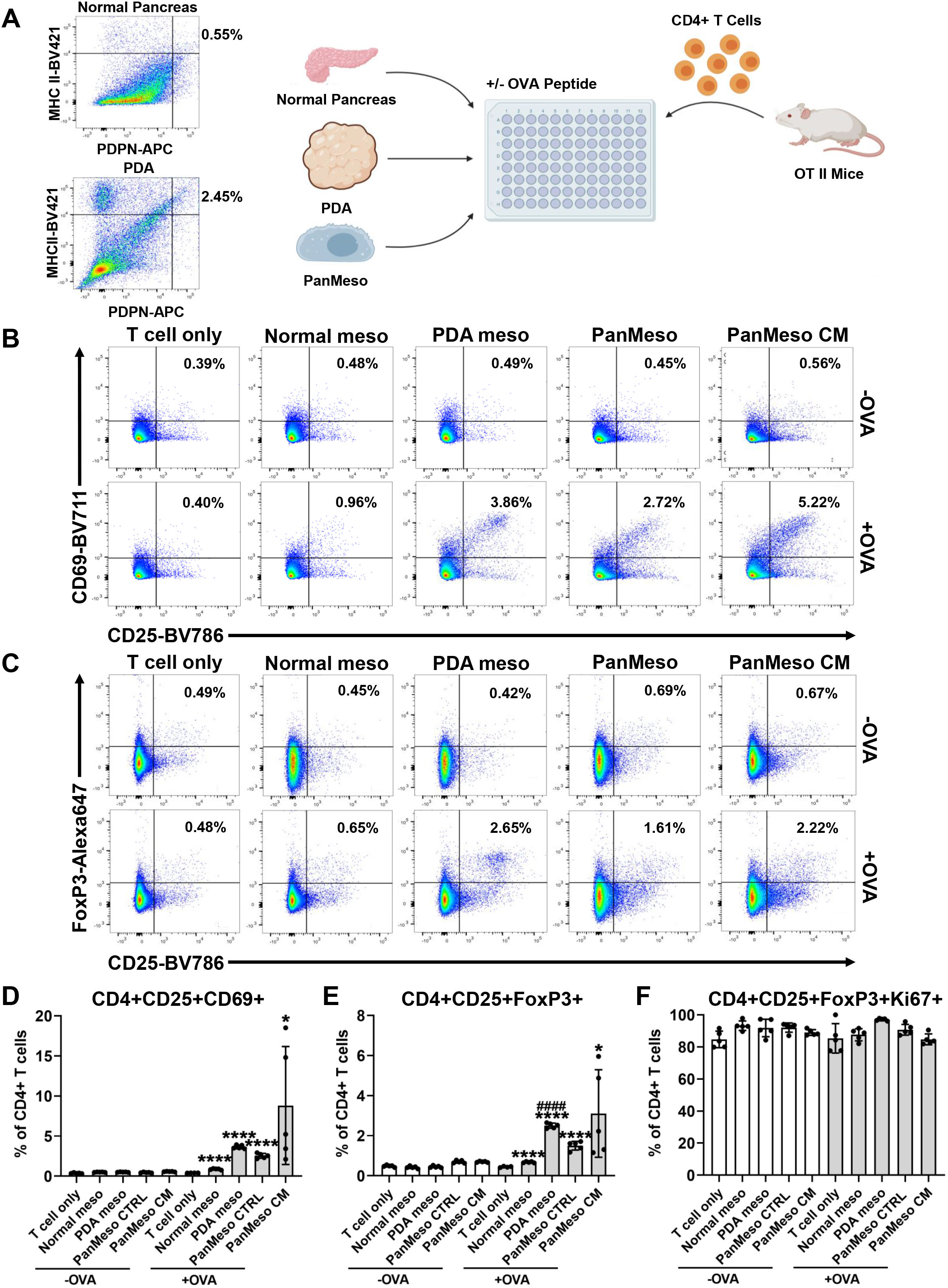
apCAFs Induce Naïve CD4^+^ T Cells into Tregs. (A) An illustration of the mesothelial cell/apCAF-T cell co-culture assay. Podoplanin^+^MHC II^+^ mesothelial cells from normal pancreata (0.55%) or apCAFs from *KPfC* tumors (2.45%) were sorted by FACS. The sorted mesothelial cells or apCAFs were incubated with the OVA peptide (OVA 323-339) for 4 hrs. Cells were then washed and co-cultured with CD4^+^ T cells isolated from OT II mice for 18 hrs. A similar assay was also performed using PanMeso cells. PanMeso cells were pre-treated with control medium or PDA organoid conditioned medium for 48 hrs and washed before the incubation with OVA peptide and being co-cultured with CD4^+^ T cells. (B-F) The CD4^+^ T cells after being co-cultured with mesothelial cells/apCAFs (A) were subjected to flow cytometry for the analysis of the early activation markers of TCR ligation CD25 and CD69 (B and D), Treg markers CD25 and FoxP3 (C and E) as well as proliferation marker Ki67 (F). *P<0.05, ****P<0.0001 vs without OVA control. ^####^ P<0.0001 vs normal mesothelial cells.

The lack of co-stimulatory molecules on professional antigen presenting cells can lead to T cell anergy or induction of regulatory T cells (Tregs) (Bour-Jordan and Bluestone, 2009; Ferrer et al., 2011; Mikami et al., 2020; Semple et al., 2011). Tregs are a subset of CD4^+^ T cells that are critical for immune tolerance. They are also a driver of immune evasion in cancer (Wing et al., 2019). Therefore, we tested the hypothesis that apCAFs induce Treg formation. We examined the presence of CD25^+^FoxP3^+^ Tregs in the CD4^+^ T cells co-cultured with OVA-loaded mesothelial cells/apCAFs. We found that mesothelial cells/apCAFs induced Treg formation in an antigen-specific manner (Figures 5C and 5E). Moreover, apCAFs had increased Treg-induction capability. To understand whether this was due to the stimulation of proliferation in pre-existing Tregs, we investigated the proliferation marker Ki67 in CD25^+^FoxP3^+^ Tregs and found that the proliferation of Treg was not increased by mesothelial cells/apCAFs (Figure 5F). This suggests that Tregs are formed through the induction by mesothelial cells/apCAFs from naïve CD4^+^ T cells directly. Taken together, these data suggest that apCAFs are an immunosuppressive CAF population that can induce Treg formation through antigen-dependent TCR ligation.

### IL-1 and TGFβ Signaling Pathways are Responsible for the Induction of apCAFs

To identify the signaling pathways that drive the mesothelial cell to apCAF transition, we extracted differential genes that were up-regulated in apCAFs and subjected them to motif enrichment analysis (Figure 6A). The gene profile of apCAFs was driven by NF-κB signaling (*Nfkb1*, *Rela*) or TGFβ signaling (*Smad1*, *Smad3*, *Smad4*). These data are highly relevant, given that these two signaling pathways have been demonstrated to be the main drivers of CAF heterogeneity in PDA (Biffi et al., 2019; Dominguez et al., 2020). NF-κB signaling, which is induced by IL-1, is responsible for the formation of the iCAF lineage, while TGFβ is responsible for the myCAF lineage during PDA progression. To validate this result and test whether these two signaling pathways were also responsible for driving mesothelial cells to differentiate into apCAFs, we treated PanMeso cells with PDA organoid CM and found that NF-κB signaling and TGFβ signaling were activated (Figure 6B). We also examined if IL-1 and TGFβ drive an apCAF phenotype in mesothelial cells. We treated PanMeso cells with IL-1, TGFβ or a combination of the two factors, and investigated the resulting gene signature by qPCR (Figures 6C and 6D). We found that either IL-1 or TGFβ could significantly down-regulate the expression of mesothelial genes (*Msln*, *Upk3b*, *Ezr*, *Nkain4*) (Figure 6C). In contrast, *Cdh11* and *Pdpn* were not down-regulated, supporting these two genes are stable markers of mesothelial cell-apCAF lineage. In comparison, we found that fibroblastic genes were differentially up-regulated by IL-1 or TGFβ (Figure 6D). For example, some genes were specifically up-regulated by IL-1 (*Cxcl1*), some were up-regulated by TGFβ (*Cxcl12*, *Col1a1*, *Col1a2*, *Col12a1*, *Tgfb1*), some could be up-regulated by either factor (*Il6*), and some depended on the presence of both factors (*Il1a*). Taken together, these qPCR data suggest that IL-1 and TGFβ can induce normal mesothelial cells into an apCAF phenotype.

**Figure 6.**
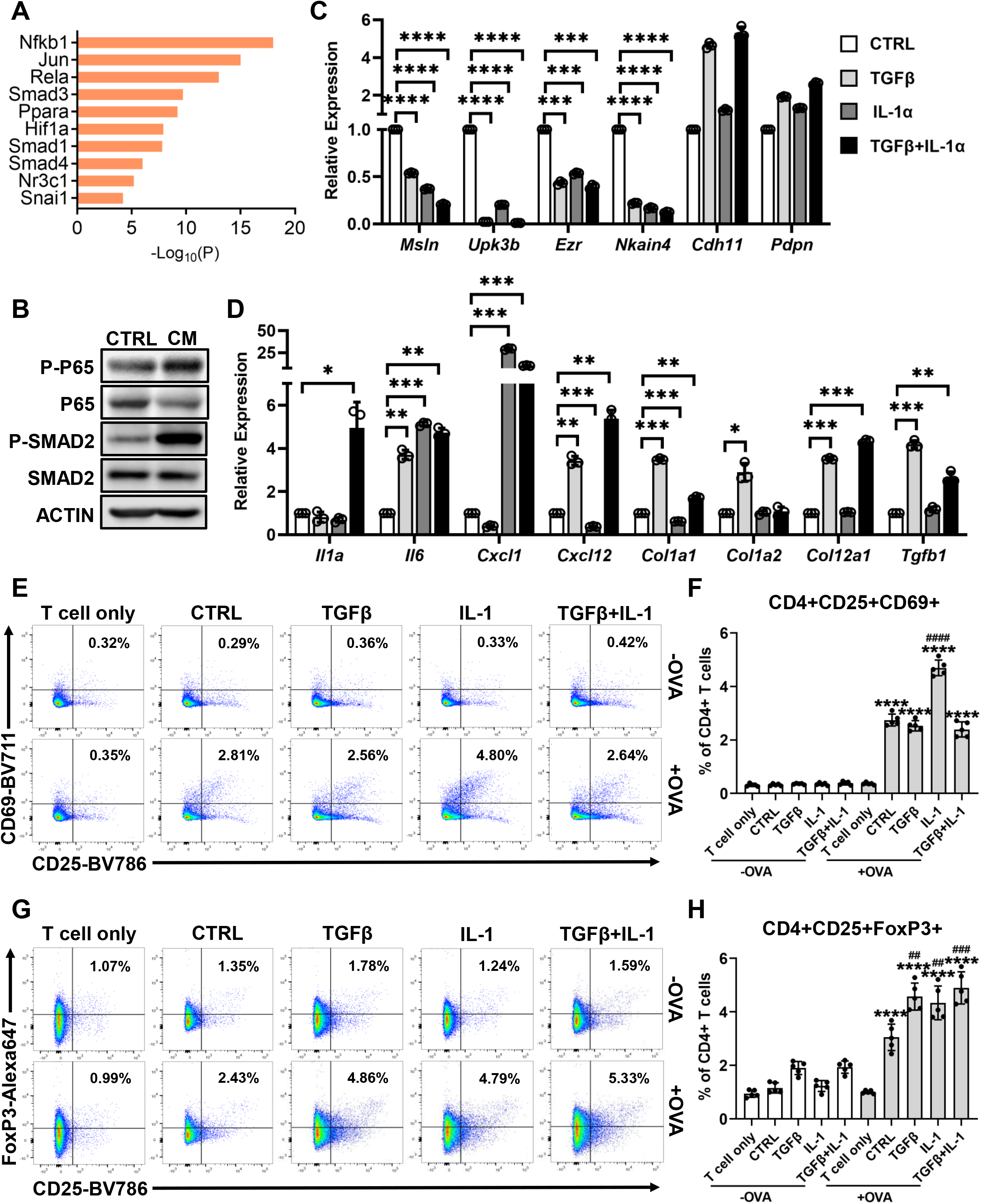
IL-1 and TGFβ are Responsible for Mesothelial Cell-apCAF Transition. (A) The differential genes that were up-regulated in apCAFs compared to normal mesothelial cells from the integrated data (Figure 1G) were subjected to motif enrichment analysis. Top transcription factors from the analysis were shown. (B) The PanMeso cells were treated with control medium or PDA organoid conditioned medium for 48 hrs. Cell lysates were harvested and subjected to western blot, probing for the phosphorylation of the NF-κB signaling (P-P65) and TGFβ signaling (P-SMAD2) proteins. (C-D) The PanMeso cells were treated with IL-1α (5 ng/mL), TGFβ (30 ng/mL) or a combination of IL-1α (5 ng/mL) and TGFβ (30 ng/mL) for 48 hrs. Cells were then harvested and subjected to qPCR for mesothelial (*Msln*, *Upk3b*, *Ezr*, *Nkain4*, *Cdh11*, *Pdpn*) (C) and fibroblastic (*Il1a*, *Il6*, *Cxcl1*, *Cxcl12*, *Col1a1*, *Col1a2*, *Col12a1*, *Tgfb1*) (D) genes. n=3, *P<0.05, **P<0.01, ***P<0.001. (E-H) The PanMeso cells were pre-treated with IL-1α (5 ng/mL), TGFβ (30 ng/mL) or a combination of IL-1α (5 ng/mL) and TGFβ (30 ng/mL) for 48 hrs and washed. Cells were then incubated with the OVA peptide (OVA 323-339) for 4 hrs. The PanMeso cells were washed and co-cultured with CD4^+^ T cells isolated from OT II mice for 18 hrs. The CD4^+^ T cells after the co-culture were subjected to flow cytometry for the analysis of the early activation markers of TCR ligation CD25 and CD69 (E and F) and the Treg markers CD25 and FoxP3 (G and H). ****P<0.0001 vs without OVA control. ^##^P<0.01, ^###^P<0.001, ^####^ P<0.0001 vs control with OVA treatment.

To examine whether IL-1 and TGFβ affect the function of mesothelial cells, we performed the mesothelial cell-CD4^+^ T cell co-culture assay using PanMeso cells (Figures 6E-6H). PanMeso cells were pre-treated with IL-1, TGFβ or a combination of the two factors for 2 days and washed before incubation with OVA peptide. CD4^+^ T cells isolated from OT II mice were then co-cultured with PanMeso cells and subjected to flow cytometry for the analysis of T cell ligation and Treg formation. We found that OVA-specific TCR activation was enhanced by pre-treatment of PanMeso cells with IL-1 but was not affected by TGFβ (Figures 6E and 6F). In contrast, IL-1 or TGFβ pre-treatment of PanMeso cells significantly enhances Treg-induction (Figures 6G and 6H). In summary, our data suggest that IL-1/NF-κB and TGFβ signaling pathways are responsible for the induction of normal mesothelial cells into apCAFs.

## Discussion

In this study, we have integrated our scRNA-seq data with other scRNA-seq datasets that have captured mesothelial cells/apCAFs in PDA. As a result, we were able to distinguish apCAFs as a unique CAF population derived from normal mesothelial cells through a fibroblastic transition. In adult vertebrates, mesothelial cells form an epithelial monolayer called mesothelium (Koopmans and Rinkevich, 2018). This layer of cells covers body cavities including thoracic and abdominal cavities as well as internal organs. Multiple lineage tracing studies have demonstrated that during embryo development, mesothelial cells are a major source of mesenchymal cells, such as fibroblasts and smooth muscle cells in developing organs and visceral white adipocytes (Chau et al., 2014; Rinkevich et al., 2012). In a recent scRNA-seq study in the developing murine pancreata, mesothelial cells were found to give rise to pancreatic fibroblastic and vascular smooth muscle cells (Byrnes et al., 2018). Under physiological conditions, the main function of adult mesothelium is to produce a lubricating fluid consisting of glycosaminoglycans that provides a slippery and non-adhesive surface to protect visceral organs and facilitate intracoelomic movement (Mutsaers, 2002). The mesothelium is also involved in the transport and movement of fluid and particles across the serosal cavities.

During organ or serosal damage, mesothelial cells can secrete pro-inflammatory chemokines and cytokines, growth factors and ECM proteins to facilitate leukocyte migration and tissue repair (Buechler et al., 2019; Mutsaers, 2002). In the recent years, studies have found that mesothelial cells are also an important source of fibroblasts during wound repair or fibrosis under many pathological circumstances, including cardiac and hepatic injury, lung fibrosis, peritoneal fibrosis and surgical adhesion (Cao and Poss, 2018; Li et al., 2013; Si et al., 2019; Sontake et al., 2018; Tsai et al., 2018). However, mesothelial cells have been traditionally neglected as a major constituent of the TME. Here for the first time, we have demonstrated that mesothelial cells can directly contribute to the TME by forming a Treg-inducing CAF population in PDA. Given that the majority of visceral organs are lined by the mesothelium, future studies are needed to determine if mesothelial cells contribute to the CAF populations in other cancer types.

Our findings are also supported by a recent study that exploited scRNA-seq to profile cell heterogeneity in patient-derived organoids (PDOs) derived from PDA biopsies (Lee et al., 2020). In this study, the three populations of CAFs (iCAFs, myCAFs and apCAFs) were found to coexist in the PDOs of primary and metastatic tumors. Moreover, the proportion of apCAFs was negatively correlated with the ratio of memory and effector CD8^+^ T cells to Tregs, highlighting the importance of apCAFs on Treg formation in human PDA. However, due to the fact that the expression of *MSLN* was not detected in apCAFs in this study, they concluded that apCAFs are not mesothelial cells. However, this is consistent with our results showing the down-regulation of mesothelial genes during mesothelial cell-apCAF transition.

In cancer, Tregs are one of the major cell types that participate in immune evasion. Treg frequency correlates with poor prognosis in PDA patients (Hiraoka et al., 2006; Tang et al., 2014). Although many studies have focused on identifying the mechanisms of Treg formation in tumors, how other stromal cells are involved in these processes and interact with Tregs is still not well understood. Multiple lines of evidence have been reported that there may be a crosstalk between CAFs and Tregs. For example, the ablation of myCAFs can increase the number of Tregs in PDA (Ozdemir et al., 2014). On the other hand, the depletion of Tregs can lead to a reduction of myCAFs (Zhang et al., 2020). Importantly, in these studies, the mesothelial cell-apCAF lineage was not examined. We predict that the mesothelial-apCAF lineage would be affected by myCAF ablation and thus the results of ablation studies should be re-examined in the context of new information defining CAF subsets more completely. Future studies are needed to better understand the relationship among different CAF populations as well as the relationship between CAFs and immune cells including Tregs.

Due to the specific location of the mesothelium, the formation of apCAFs likely occurs at the periphery of the tumor. This is highly relevant, given that the periphery is considered to be an area where blood vessels enter and exit the tumor. Therefore, it is tempting to speculate that tumor mesothelium may create the first stromal barrier that immune cells encounter and this way contributes to the immune evasive phenotype, such as T cell exclusion (Jerby-Arnon et al., 2018). Further studies are required to elucidate and define the contribution of mesothelial cells/apCAFs to the regional heterogeneity of cancer cells and TME. Given that mesothelial cells can express iCAF and myCAF genes during the transition to apCAFs, this adds additional complexity to the specificity of current iCAF and myCAF markers. Therefore, the identification of a specific CAF population may require multiple markers or a transcriptomic approach. Although we have identified IL-1 and TGFβ as two signals that can induce a fibroblastic phenotype and Treg-inducing capability in mesothelial cells, further investigation is needed to specifically target the signaling pathways that mediate the mesothelial cell-apCAF transition and determine the therapeutic potential of such targeted therapies.

In summary, we have identified mesothelial cells as the origin of apCAFs. Through the fibroblastic transition, mesothelial cells contribute to the formation of an important Treg-inducing CAF population. This might be an important mechanism of immune evasion in cancer. Future studies need to further understand this biology, which will potentially lead to the identification of novel strategies to target CAFs and overcome immune evasion in PDA as well as other cancer types.

## Supporting information

Key Resources Table

## Acknowledgements

This work was supported by NIH grants R01 (CA243577) and U54 (CA210181 Project 2), the Effie Marie Cain Fellowship, and the Jean Shelby Fund for Cancer Research at Communities Foundation of Texas to RAB. We acknowledge helpful input from other members of the Brekken laboratory and Drs. Michael Dellinger, Eric N. Olson and Rana Gupta. We thank Dr. Shannon Turley for sharing information of their scRNA-seq study. We also thank UT Southwestern Core facilities supported in part by the Cancer Center Support Grant (P30 CA142543), core facilities used included the McDermott Center Next-Generation Sequencing Core, the Live Cell Imaging Core and the Flow Cytometry Core.

## Author Contributions

HH: study concept and design, acquisition of data, analysis, interpretation of data and drafting of the manuscript. ZW: acquisition of data and bioinformatic analyses. YZ: acquisition of data. RAB: study concept and design, interpretation of data, and drafting of the manuscript.

## STAR Methods

### Resource Availability

#### Lead Contact

Further information and requests for resources should be directed to and will be fulfilled by the Lead Contact, Huocong Huang

### Materials Availability

All unique reagents generated in this study are available from the Lead Contact with a completed Materials Transfer Agreement.

### Data and Code Availability

This study did not generate new raw scRNA-seq data. The raw scRNA-seq data we used to extend our analyses on fibroblasts in this study have been previously uploaded to the National Center for Biotechnology Information’s Gene Expression Omnibus database repository (https://www.ncbi.nlm.nih.gov/geo) under accession number GSE125588.

### Experimental Model and Subject Details

#### Cell Lines

BMFA3 and CT1BA5 were mouse primary pancreatic cancer cell lines derived from *KPfC* mice on a C57BL/6 background and isolated as described previously (Huang et al., 2019; Ludwig et al., 2018). PanMeso cells were established as follows. Mesothelium was harvested from normal pancreata of immortomice under the dissecting microscope. The tissues of immortomice carry the transgene that allows the expression of a large tumor antigen at the permissive temperature of 33 °C in the presence of IFN-γ (Jat et al., 1991), while the large tumor antigen would be rapidly denatured at the nonpermissive temperature of 37 °C. The pancreatic mesothelium was seeded onto a tissue culture dish and cultured at 33 °C. Cells were subjected to FACS when they had been cultured for 20 days and became confluent. Podoplanin^+^MHC II^+^ cells were collected and named pancreatic mesothelial (PanMeso) cells. All mouse primary pancreatic cancer cell lines were cultured in DMEM (Invitrogen) containing 10% FBS and 1% penicillin/streptomycin (Gibco/Thermo) and maintained at 37°C in an atmosphere of 5% CO2. The PanMeso cells were cultured in the mesothelial cell media (medium 199 (Gibco/Thermo), 10% FBS, 1% penicillin/streptomycin (Gibco/Thermo), 3.3 nM mouse epidermal growth factor (Biolegend), 400 nM hydrocortisone (MilliporeSigma), 870 nM zinc-free bovine insulin (MilliporeSigma), 20 mM HEPES (Thermo Fisher Scientific)). For maintenance, the PanMeso cells were cultured at 33°C with 10 units/ml IFN-γ (R&D Systems) in an atmosphere of 5% CO2. For experiments, the cells were washed with PBS three times and cultured in mesothelial cell media without IFN-γ at 37°C in an atmosphere of 5% CO2 for 5 days before any experiments. All cell lines in this study were confirmed to be free of mycoplasma (e-Myco kit, Boca Scientific) before use.

#### Animals

*KIC* and *KPfC* mice were generated as previously described (Hingorani et al., 2003; Hingorani et al., 2005). Mice were sacrificed when they had early stage tumors (40-day-old) or late stage (60-day-old) tumors for *KIC* and *KPfC* mice. The *KIC* mice are on a mixed background (C57BL/6 with FVB). The *KPfC* mice are a pure C57BL/6 genetic background. For staining and lineage tracing assays, at least 6 mice (male and female) were used per time point. Normal pancreata were obtained from Cre-negative littermates of the *KIC* or *KPfC* mice. 6-week-old mice male and female mice were purchased from The Jackson Laboratory, including C57BL/6J mice C57Bl/6J (JAX stock #000664), immortomice (JAX stock #032619) and OT II mice (JAX stock #004194). All animals were housed in a pathogen-free facility with 24-hr access to food and water. Animal experiments in this study were approved by and performed in accordance with the institutional animal care and use committee at the UTSW Medical Center at Dallas. Mice were euthanized by cervical dislocation under anesthesia.

### Method Details

#### Plasmids

pLV-eGFP (Addgene Plasmid #36083) was used to expressed eGFP in PanMeso cells.

#### Immunohistochemistry

Formalin-fixed, paraffin-embedded tissues were cut in 5-μm sections. Sections were evaluated by single color or multiplex immunohistochemical analysis following our previously reported protocol (Sorrelle et al., 2019) using antibodies specific for cadherin-11 (LSBio, 1:300), podoplanin (MilliporeSigma, 1:300), mesothelin (LSBio, 1:1000), CD74 (Biolegend, 1:300), α-SMA (Biocare Medical, 1:500) or eGPF (Abcam, 1:200). For brightfield images, slides were scanned by NanoZoomer 2.0-HT digital slide scanner (Hamamatsu) and images were obtained with NDP.view2 software (Hamamatsu). For fluorescent images, images were obtained and analyzed with the Zeiss LSM 780 confocal microscope and ZEN software (Zeiss).

#### RT-PCR

RNA was extracted using RNeasy Mini Kit (Qiagen). cDNAs were synthesized from 1 μg of total RNA using the iScript cDNA synthesis kit (Bio-Rad). The expression of genes was measured by qRT-PCR with SYBR-Green Master Mix (Bio-Rad). For a detailed list of qRT-PCR primer sequences, see Table S1.

#### Western Blot Analysis

Cells were extracted with radioimmunoprecipitation assay buffer (50 mM Tris-HCl, pH 8.0, 150 mM NaCl, 0.1% SDS, 0.5% sodium deoxycholate, and 1% Nonidet P-40) for SDS-PAGE. Protein concentration was determined using a BCA Protein Assay Kit (Thermo Fisher Scientific). Laemmli sample buffer was then added to the protein lysates and boiled for 10 mins. The proteins were resolved by SDS-PAGE, electrophoretically transferred to nitrocellulose membranes, and blocked in 5% nonfat dry milk or bovine serum albumin. Blocking buffer was then removed, and the membranes were incubated with primary antibody in TBST (10 mM Tris-HCl, pH 7.5, 150 mM NaCl, 0.05% Tween 20) for 1 hr, then washed 3 × 10 mins with TBST, and incubated with horseradish peroxidase-conjugated anti-mouse or rabbit secondary antibody (Jackson ImmunoResearch Laboratories) for 1 hr. The secondary antibody was removed by washing 3 × 10 mins with TBST. The membranes were incubated with SuperSignal West Pico substrate (Thermo Fisher Scientific) for the detection of the immunoreactive bands.

#### Orthotopic Models

BMFA3 or CT1BA5 cells were injected orthotopically (5 × 10^5^ cells) with or without PanMeso cells (5 × 10^5^ cells) in 6-week-old C57BL/6 mice (male and female, n=6/group). To harvest early stage tumors, mice were sacrificed 1 week after injection. To harvest late stage tumors, mice were sacrificed 1 month after injection. Tumors were then subjected to IHC staining.

#### Lineage-tracing Assay

CellTrace CFSE dye (Thermo Fisher Scientific, 1:1000, 2mL/mouse) was injected intraperitoneally into C57BL/6 mice bearing normal pancreata or late-stage-tumor-bearing *KIC* or *KPfC* mice (60-day-old). Normal pancreata or PDAs were harvested 2 days after the injection, snap freezed by liquid nitrogen, and cut into 10-μm frozen sections. Tissues were then fixed by ice-cold 100% methanol for 15 mins at −20°C and rinsed 3 × 5 mins with PBS (1.86 mM NaH2PO4, 8.41 mM Na2HPO4, 175mM NaCl, pH 7.4). For CFSE signal imaging, fixed tissues were directly coverslipped with Prolong Gold Antifade Reagent with DAPI (Cell Signaling). For co-staining, fixed tissues were then blocked with 5% bovine serum albumin in PBST (1.86 mM NaH_2_PO_4_, 8.41 mM Na_2_HPO_4_, 175mM NaCl, 0.05% Tween 20, pH 7.4) for 1 hr, incubated with diluted primary antibody against cadherin-11 (LSBio, 1:200), SOX9 (MilliporeSigma, 1:200) or α-SMA (Biocare Medical, 1:500) for 1hr, rinsed 3 × 5 mins with PBST, incubated with diluted secondary antibody for 1hr, then rinsed 3 × 5 mins with PBST and coverslipped with Prolong Gold Antifade Reagent with DAPI. Images were obtained and analyzed with the Zeiss LSM 780 confocal microscope and ZEN software (Zeiss).

#### Tumor Organoid Conditioned Medium

Mouse PDA organoids were isolated from *KPfC* mice with late stage tumors. Tumor tissues were minced and digested at 37°C for 12 hr in a digestion buffer containing 0.012% collagenase XI (MilliporeSigma) and 0.012% dispase (Gibco) in DMEM (Invitrogen) containing 1% FBS. The dissociated organoids were then pelleted, washed with PBS and seeded on petri dish in DMEM (Invitrogen) containing 10% FBS. After 4 days of culture, the medium was filtered through a 0.22 μm ultra-low protein binding filter and collected as conditioned medium.

#### Sorting of Mesothelial Cells or apCAFs

The digestion buffer was prepared in DMEM with 1% FBS: collagenase type I (45 units/mL, Worthington Biochemical), collagenase type II (15 units/mL, Worthington Biochemical), collagenase type III (45 units/mL, Worthington Biochemical), collagenase type IV (45 units/mL, Gibco, Thermo Fisher Scientific), elastase (0.08 units/mL, Worthington Biochemical), hyaluronidase (30 units/mL, MilliporeSigma), and DNAse type I (25 units/mL, MilliporeSigma). Normal pancreata or tumors were enzymatically digested into a single cell suspension as follow. Freshly dissected tissues were minced in a 10 cm petri dish. Samples were then resuspended in 10mL digestion buffer and incubated on a shaker at 37°C for 60 mins. Then 30 mL of DMEM containing 1% FBS was added, and cells were washed 3 times with PBS before filtering through a 70-μm mesh filter (MilliporeSigma). Single cells were resuspended in PBS with 1% FBS, blocked with anti-mouse CD16/CD32 Fc Block for 15 mins at 4°C (Biolegend, clone 2.4G2, 1:50) and then incubated with antibodies against podoplanin (Biolegend, 1:200) and I-A/I-E (Biolegend, MHC class II, 1:200) for 30 mins at 4°C. Cells were washed with PBS containing 1% FBS 3 times and subjected to cell sorting on the FACSAria sorter (UTSW flow cytometry core). Sorted cells were collected into ice cold DMEM with 10% FBS.

#### Mesothelial Cell/apCAF-CD4+ T cell Coculture Assay and Flow Cytometry Analysis

20,000 sorted normal mesothelial cells/apCAFs or PanMeso cells were seeded in U-bottom 96-well plates and incubated with or without 25 μg/ml OVA peptide (OVA 323-339) (GenScript) in DMEM with 10% FBS for 4 hrs at 37°C. Spleens from 6-to-8-week-old OT II mice were harvested and CD4^+^ T cells were isolated using a MojoSort mouse CD4^+^ T cell isolation kit (Biolegend). Mesothelial cells/apCAFs were washed 3 times and co-cultured with 50,000 CD4^+^ T cells in DMEM with 10% FBS for 18 hrs. T cells were then washed twice with PBS and stained with Fixable Viability Stain (BD Biosciences) for 1 hour. The cell suspensions were then washed and stained with antibodies against CD4 (BD Biosciences, 1:100), CD25 (BD Biosciences, 1:100) and CD69 (BD Biosciences, 1:100) for 1 hr at 4°C. Surface-stained cells were fixed, permeabilized, and stained for intracellular markers FoxP3 (BD Biosciences, 1:100) and Ki67 (Biolegend, 1:100). Cells were analyzed using FACS LSRFortessa SORP, and the FlowJo software.

#### scRNA-seq Data Preprocessing

The scRNA-seq data were analyzed using R package Seurat v3.2.2 (Butler et al., 2018; Stuart et al., 2019). Fibroblasts scRNA-seq data from Hosein et al. were analyzed as previously described (Hosein et al., 2019), and converted to Seurat v3 format using the function UpdateSeuratObject. The scRNA-seq data by Dominguez et al. were obtained from the ArrayExpress database under the accession number E-MTAB-8483, and quality control filtering was performed as described previously (Dominguez et al., 2020). Cell clusters were identified using the first 12 principle components under the clustering resolution of 0.6. Identities of the cell clusters were matched to the previously reported cluster names based on the expression of genes mentioned in Dominguez et al. (Dominguez et al., 2020). The scRNA-seq data by Elyada et al. were obtained from the Gene Expression Omnibus under the accession number GSE129455 (Elyada et al., 2019). Identities of the cell clusters were matched to the previously reported cluster names based on the expression of genes mentioned in Elyada et al. (Elyada et al., 2019).

#### Integration of Multiple scRNA-seq Datasets

The datasets by Hosein et al., Dominguez et al., and Elyada et al. (Dominguez et al., 2020; Elyada et al., 2019; Hosein et al., 2019) were integrated as described in Seurat v3.2 Vignettes using the Standard Workflow (https://satijalab.org/seurat/v3.2/integration.html). Briefly, top 2,000 variable features were identified for each dataset. Next, integration anchors were identified using the first 30 dimensions, and the datasets were integrated using Seurat function IntegrateData. The integrated dataset was subsequently rescaled, principle component analysis was performed, and dimensional reduction was performed using the first 20 principle components. The cell clusters were identified from the integrated dataset using the first 12 principle components under the clustering resolution of 0.6. Differential expression analysis was performed using the Seurat functions FindMarkers and FindAllMarkers.

#### Pseudotime Analysis

R package Monocle2 (v2.16.0) was used to align cells from clusters 1, 4, 7, 8, 9, 10 of the integrated dataset (Qiu et al., 2017). The cells from Elyada et al. (Elyada et al., 2019) were not included in this analysis due to the lack of raw read counts. The top 1,000 differentially expressed genes were used to order cells in the pseudotime trajectories, determined using the Monocle2 function differentialGeneTest. Dimensional reduction was performed by the Monocle2 function reduceDimension using DDRTree method.

#### Gene Ontology Analysis and Motif Enrichment Analysis

Gene ontology analysis and motif enrichment analysis were performed using Metascape (Zhou et al., 2019). The motif enrichment analysis from Metascape was based on the TRRUST algorithm.

#### Quantification and Statistical Analysis

All purchased mice in this study had similar age. Male and female mice were included in equal numbers for each animal experiment. All data are reported as mean ± SD. Statistical analysis was performed with a 2-tailed t-test using GraphPad Prism software and R language (https://www.R-project.org/). P < 0.05 was considered statistically significant. All in vitro experiments were performed with at least three biological replicates. Illustrations were created with BioRender.com.

