## Supplementary material for "Mesothelial cell-derived antigen-presenting cancer-associated fibroblasts induce expansion of regulatory T cells in pancreatic cancer": Key Resources Table

| REAGENT or RESOURCE | SOURCE | IDENTIFIER |
|---|---|---|
| Antibodies | | |
| Chicken Anti-GFP | Abcam | Cat# ab13970 |
| BV421 Rat Anti-CD4 (clone GK1.5) | BD Biosciences | Cat# 562891 |
| BV786 Rat Anti-CD25 (clone PC61) | BD Biosciences | Cat# 564023 |
| BV711 Hamster Anti-CD69 (Clone H1.2F3) | BD Biosciences | Cat# 740664 |
| Alexa Fluor 647 Rat anti-Foxp3 (Clone MF23) | BD Biosciences | Cat# 560401 |
| Mouse Anti-Smooth Muscle Actin (1A4) | Biocare Medical | Cat# 001 |
| Rat Anti-CD74 (CLIP) | Biolegend | Cat# 151002 |
| Brilliant Violet 421 Rat Anti-I-A/I-E (M5/114.15.2) | Biolegend | Cat# 107632 |
| PE Anti-Ki-67 (16A8) | Biolegend | Cat# 652404 |
| APC Anti-Podoplanin (clone 8.1.1) | Biolegend | Cat# 127410 |
| Rabbit Phospho-NF-κB p65 (Ser536) (93H1) | Cell Signaling | Cat# 3033S |
| Mouse Anti-Smad2 (L16D3) | Cell Signaling | Cat# 3103S |
| Rabbit Anti-Phospho-Smad2 (Ser465/467) (138D4) | Cell Signaling | Cat# 3108S |
| Peroxidase AffiniPure Donkey Anti-Chicken IgY (IgG) (H+L) | Jackson ImmunoResearch | Cat# 703-035-155 |
| Rabbit Anti-Cadherin-11 | LSBio | Cat# LS-B2308 |
| Rabbit Anti-Mesothelin | LSBio | Cat# LS-C407883 |
| Rabbit Anti-Actin | MilliporeSigma | Cat# A2066 |
| Syrian Hamster Anti-Podoplanin (clone 8.1.1) | MilliporeSigma | Cat# MABT1512 |
| Rabbit Anti-Sox9 | MilliporeSigma | Cat# AB5535 |
| Rabbit Anti-NFκB p65 | Santa Cruz | Cat# sc-109 |
| Goat anti-Mouse IgG (H+L) Highly Cross-Adsorbed Secondary Antibody, Alexa Fluor 546 | Thermo Fisher Scientific | Cat# A-11030 |
| Goat anti-Rabbit IgG (H+L) Highly Cross-Adsorbed Secondary Antibody, Alexa Fluor 546 | Thermo Fisher Scientific | Cat# A-11035 |

| | | |
|---|---|---|
| Bacterial and Virus Strains | | |
| pLV-eGFP | Addgene | Cat# 36083 |
| Chemicals, Peptides, and Recombinant Proteins | | |
| OVA Peptide (323-339) | GenScript | Cat# RP10610 |
| Recombinant Human TGF-β1 (CHO derived) | PeproTech | Cat# 100-21C |
| Recombinant Mouse IL-1 alpha/IL-1F1 Protein | R&D Systems | Cat# 400-ML-005/CF |
| Critical Commercial Assays | | |
| Fixable Viability Stain | BD Biosciences | Cat# 564406 |
| MojoSort mouse CD4 T cell isolation kit | Biolegend | Cat# 480033 |
| iScript cDNA Synthesis Kit | Bio-Rad | Cat# 1708891 |
| iTaq Universal SYBR® Green Supermix | Bio-Rad | Cat# 1725121 |
| Opal 520 Reagent Pack | PerkinElmer | Cat# FP1487001KT |
| Opal 570 Reagent Pack | PerkinElmer | Cat# FP1488001KT |
| RNeasy Mini Kit | Qiagen | Cat# 74106 |
| CellTrace CFSE Cell Proliferation Kit | Thermo Fisher Scientific | Cat# C34554 |
| ImmPRESS AP Horse Anti-Rabbit IgG Polymer Detection Kit | Vector Laboratories | Cat# MP-5401-50 |
| ImmPRESS AP Horse Anti-Mouse IgG Polymer Detection Kit | Vector Laboratories | Cat# MP-5402-50 |
| ImmPRESS HRP Goat Anti-Rabbit IgG Polymer Detection Kit | Vector Laboratories | Cat# MP-7451-50 |
| Experimental Models: Cell Lines | | |
| Mouse Cell Line: BMFA3 | Huang et al., 2019 | N/A |
| Mouse Cell Line: CT1BA5 | Huang et al., 2019 | N/A |
| Mouse Cell Line: PanMeso | This Paper | N/A |
| Experimental Models: Organisms/Strains | | |
| Mouse: *KIC* ($Kras^{LSL-G12D/+}$;$Ink4a^{fl/fl}$;$Ptf1a^{Cre/+}$) | Hingorani et al., 2005 | N/A |

| | | |
|---|---|---|
| Mouse: *KPfC* (*Kras*$^{LSL-G12D/+}$;*Trp53*$^{fl/fl}$;*Pdx1*$^{Cre/+}$) | Hingorani et al., 2003 | N/A |
| Mouse: C57BL/6J | The Jackson Laboratory | Cat# 000664 |
| Mouse: Immortomouse (Tg(H2-K1-tsA58)6Kio/LicrmJ | The Jackson Laboratory | Cat# 032619 |
| Mouse: OT II (B6.Cg-Tg(TcraTcrb)425Cbn/J) | The Jackson Laboratory | Cat# 004194 |
| Software and Algorithms | | |
| R package Seurat (v3.2.2) | Butler et al., 2018<br>Stuart et al., 2019 | N/A |
| R package Monocle2 (v2.16.0) | Qiu et al., 2017 | N/A |
| R language | R-project | https://www.R-project.org/ |
| FlowJo software (v10.7) | BD Biosciences | https://www.flowjo.com/ |
| Graphpad Prism 8.0 software | GraphPad Software, Inc. | http://www.graphpad.com/scientific-software/prism/ |
| ZEN Imaging Software | Zeiss | https://www.zeiss.com/microscopy/us/products/microscope-software/zen.html |
| NDP.view2 software | Hamamatsu Photonics | https://www.hamamatsu.com/us/en/product/type/U12388-01/index.html |
| Metascape | Zhou et al., 2019 | https://metascape.org/ |
