## Supplemental Legends for "Mesothelial cell-derived antigen-presenting cancer-associated fibroblasts induce expansion of regulatory T cells in pancreatic cancer"

### Supplemental Information

### Supplementary Figure Legends

**Figure S1. Preprocessing of the scRNA-seq Datasets for the Integrated Analyses, Related to Figure 1.**

(A) scRNA-seq data of the *KIC* PDA by Dominguez et al. were reprocessed. Identities of the cell clusters were matched to the previously reported cluster names based on the expression of genes mentioned in Dominguez et al. including clusters 0, 1, 2, 3, 4, 8, proliferating fibroblasts (Prolif.FB), normal mesothelial cells (Meso), fEMT tumor cells (fEMT), pEMT tumor cells (pEMT), myeloid cells, acinar cells and endothelial cells (EC).

(B) The heatmap displaying the top marker genes for each cell cluster of the reprocessed data from Dominguez et al. These marker genes were also used to define these cell clusters in Dominguez et al. The heatmap demonstrated our reprocessed data was consistent with Dominguez et al.

(C) scRNA-seq data of the *KPC* PDA by Elyada et al. were reprocessed. Identities of the cell clusters were matched to the previously reported cluster names based on the expression of genes mentioned in Elyada et al. including iCAFs, apCAFs and myCAFs.

(D) The heatmap displaying the top marker genes for each cell cluster of the reprocessed data from Elyada et al. These marker genes were also used to define these cell clusters in Elyada et al. The heatmap demonstrated our reprocessed data was consistent with Elyada et al.

**Figure S2. Marker Genes of Each cell cluster in the Integrated Data of Fibroblasts, Related to Figure 1.**

Three scRNA-seq datasets of fibroblasts in PDA (Hosein et al., Dominguez et al. and Elyada et al.) were integrated. Graph-based clustering of cells with UMAP was performed with the integrated data and 11 clusters of fibroblasts were identified (Figure 1G). The top marker genes for each cell cluster of the integrated data were shown as the heatmap.
