## Supplementary figures and images for "Mesothelial cell-derived antigen-presenting cancer-associated fibroblasts induce expansion of regulatory T cells in pancreatic cancer"

### Supplemental Figures

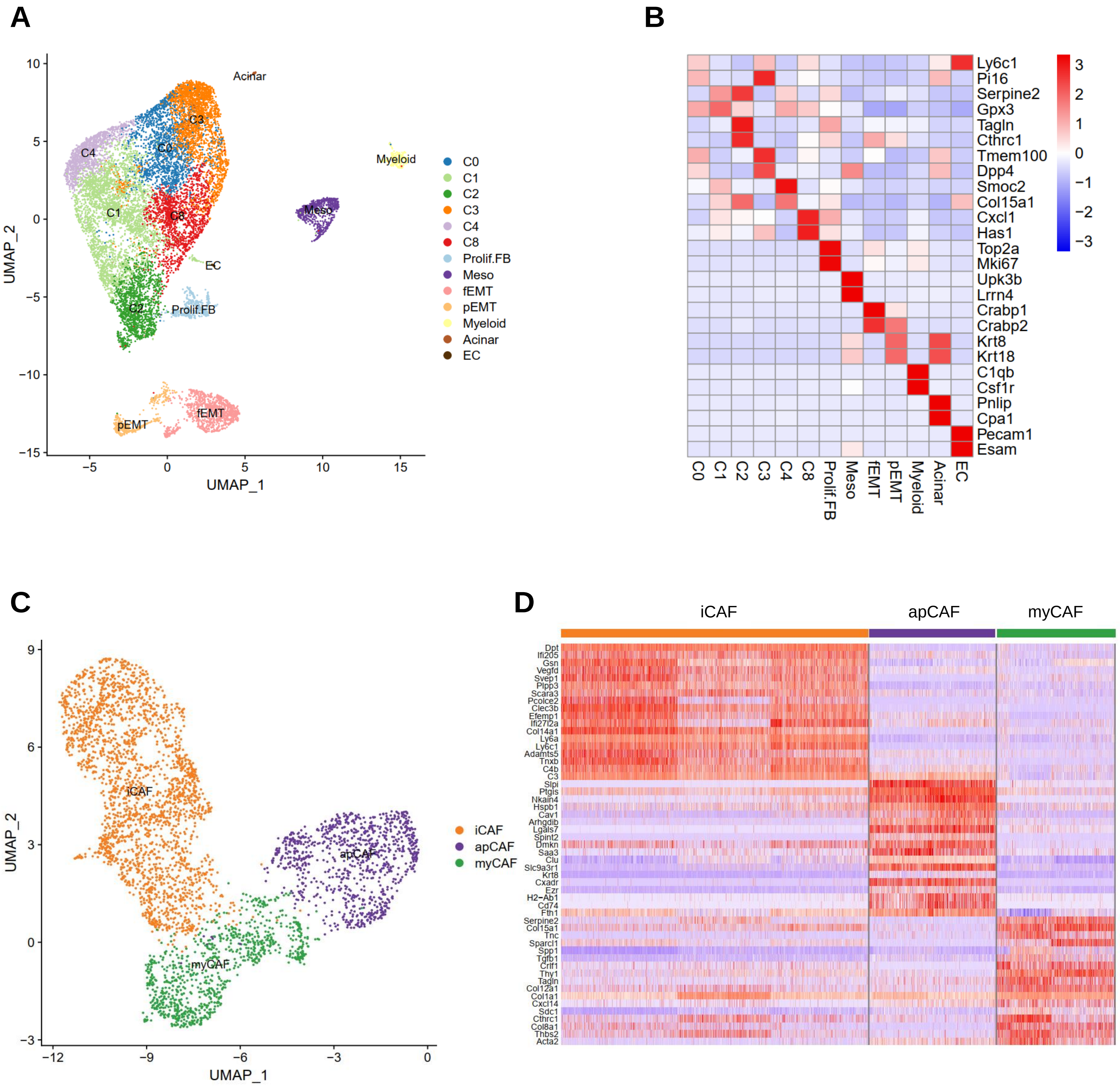

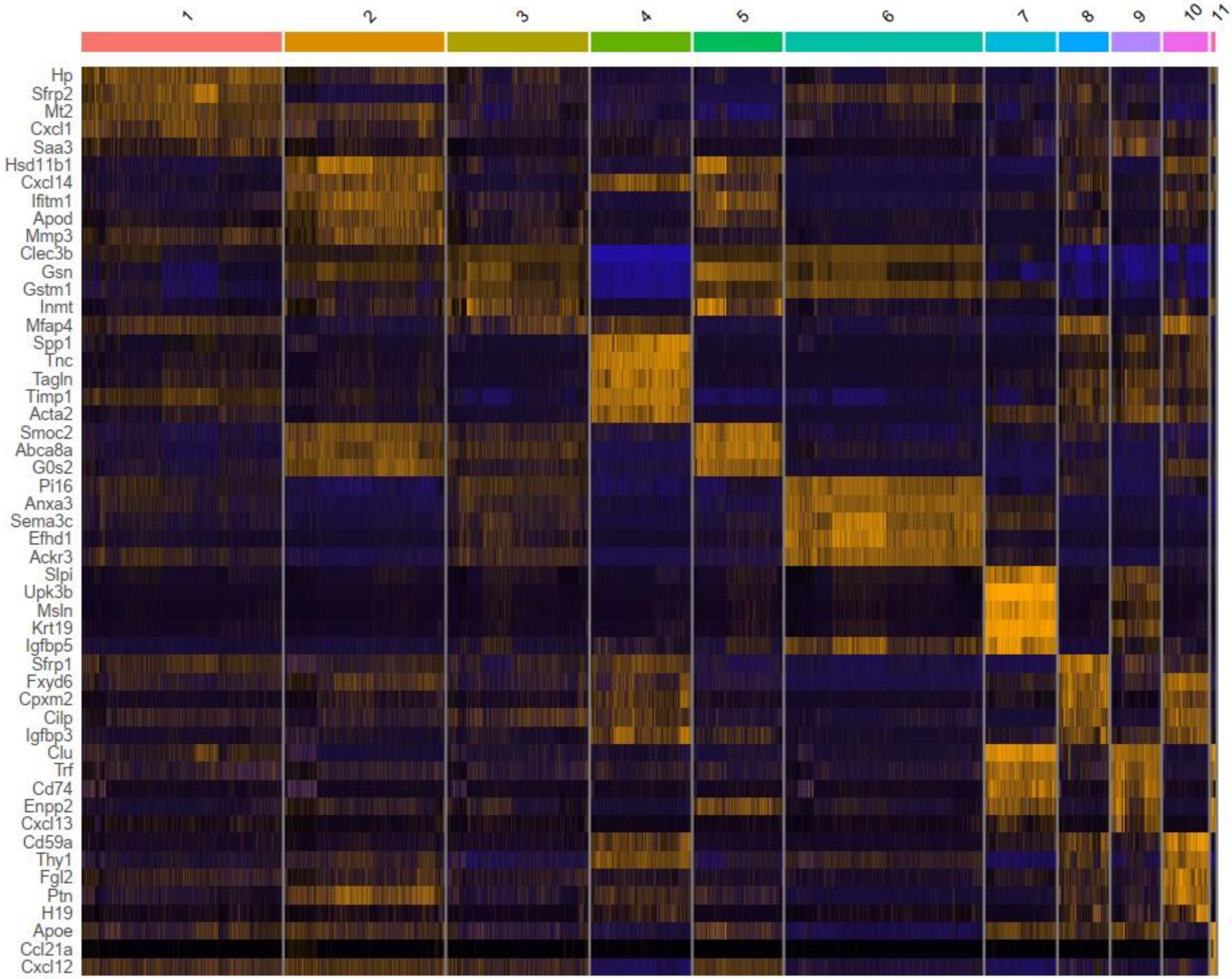
