## Supplemental Table for "Mesothelial cell-derived antigen-presenting cancer-associated fibroblasts induce expansion of regulatory T cells in pancreatic cancer"

### Table S1. qRT-PCR primer sequences

| Mouse Gene Name | Primer Sequence |
|---|---|
| *Msln* | Forward Primer: CTTGGGTGGATACCACGTCTG<br>Reverse Primer: CTTCTGTCTTACAGCCATAGCC |
| *Upk3b* | Forward Primer: AGACCTGATTGCCTACGTGC<br>Reverse Primer: GGTGTCCTTAGTTGAGACATGCT |
| *Ezr* | Forward Primer: CAATCAACGTCCGGGTGAC<br>Reverse Primer: GCCAATCGTCTTTACCACCTGA |
| *Nkain4* | Forward Primer: CTCTGGAACGGCAAGTCTTTG<br>Reverse Primer: GTGGCCGGTATTGAATGGTG |
| *Pdpn* | Forward Primer: ACCGTGCCAGTGTTGTTCTG<br>Reverse Primer: AGCACCTGTGGTTGTTATTTTGT |
| *Cdh11* | Forward Primer: CTGGGTCTGGAACCAATTCTTT<br>Reverse Primer: GCCTGAGCCATCAGTGTGTA |
| *Cd74* | Forward Primer: AGTGCGACGAGAACGGTAAC<br>Reverse Primer: CGTTGGGGAACACACACCA |
| *H2-Ab1* | Forward Primer: AGCCCCATCACTGTGGAGT<br>Reverse Primer: GATGCCGCTCAACATCTTGC |
| *Cdh2* | Forward Primer: AGCGCAGTCTTACCGAAGG<br>Reverse Primer: TCGCTGCTTTCATACTGAACTTT |
| *Vim* | Forward Primer: CGTCCACACGCACCTACAG<br>Reverse Primer: GGGGGATGAGGAATAGAGGCT |
| *Snai1* | Forward Primer: CACACGCTGCCTTGTGTCT<br>Reverse Primer: GGTCAGCAAAAGCACGGTT |
| *Snai2* | Forward Primer: TGGTCAAGAAACATTTCAACGCC<br>Reverse Primer: GGTGAGGATCTCTGGTTTTGGTA |
| *Zeb1* | Forward Primer: GCTGGCAAGACAACGTGAAAG<br>Reverse Primer: GCCTCAGGATAAATGACGGC |
| *Il6* | Forward Primer: TAGTCCTTCCTACCCCAATTTCC<br>Reverse Primer: TTGGTCCTTAGCCACTCCTTC |
| *Cxcl1* | Forward Primer: CTGGGATTCACCTCAAGAACATC<br>Reverse Primer: CAGGGTCAAGGCAAGCCTC |
| *Pdgfrb* | Forward Primer: TTCCAGGAGTGATACCAGCTT<br>Reverse Primer: AGGGGGCGTGATGACTAGG |
| *Tagln2* | Forward Primer: CCTGGCCGTGAGAACTTCC<br>Reverse Primer: GTCCGTGGTGTTAATGCCATAG |
| *Col1a1* | Forward Primer: GCTCCTCTTAGGGGCCACT<br>Reverse Primer: CCACGTCTCACCATTGGGG |
| *Col12a1* | Forward Primer: AAGTTGACCCACCTTCCGAC<br>Reverse Primer: GGTCCACTGTTATTCTGTAACCC |
| *Tgfb1* | Forward Primer: CTCCCGTGGCTTCTAGTGC<br>Reverse Primer: GCCTTAGTTTGGACAGGATCTG |
| *Il1a* | Forward Primer: CGAAGACTACAGTTCTGCCATT<br>Reverse Primer: GACGTTTCAGAGGTTCTCAGAG |
| *Cxcl12* | Forward Primer: TGCATCAGTGACGGTAAACCA<br>Reverse Primer: TTCTTCAGCCGTGCAACAATC |
| *Col1a2* | Forward Primer: GTAACTTCGTGCCTAGCAACA<br>Reverse Primer: CCTTTGTCAGAATACTGAGCAGC |
